# Knowledge-Driven Interpretable Neural Networks for Mechanistic Insight

**DOI:** 10.1101/2025.09.08.674894

**Authors:** Yan Ke, Tianwei Yu

## Abstract

Analyzing omics data based on pathway knowledge is critical for understanding the molecular mechanism behind pathological changes, but current pathway analysis methods do not model the detailed mechanistic nature of biological interactions, limiting the understanding of pathway behavior to a relatively shallow level. To address this issue, we present a knowledge-driven machine learning framework that embeds features into pathway graphs and models reactions analytically, producing interpretable feature hierarchies and sub-networks where functional associations are estimated to model biological interactions. The approach is agnostic to feature selection, enabling the use of full omics datasets without discarding weak signals. Applications to breast cancer microRNA-gene regulation data and COVID-19 metabolomics data highlight immune and metabolic pathways relevant to disease progression. This framework bridges predictive modeling with mechanistic interpretation, offering a foundation for integrative pathway analysis.

## 1 Introduction

Elucidating the mechanistic basis of biological pathways and their response to pathological changes is a foundational challenge in systems biology and clinical research. As the functional units of cellular processes, pathways orchestrate complex interactions among genes, proteins, and metabolites, thereby influencing the progression of diseases and the response to therapeutic interventions [1]. The proliferation of high-throughput multi-omics technologies has produced copious data depicting the behavior of these molecular entities at single-feature resolution, facilitating the identification of biomarkers and network markers associated with disease states [2, 3].

However, the mechanistic interpretation of these complex datasets remains a central challenge. Traditional approaches that focus on single-gene or feature analysis often miss the larger picture of coordinated, functional biological processes; such reductionist methods frequently yield lists of putative biomarkers that are difficult to validate mechanistically and clinically[4]. Pathway-level analysis has emerged as a powerful alternative, allowing researchers to contextualize molecular changes within functional networks, improve statistical power, reduce false positives, and produce biologically interpretable results. This paradigm shift has accelerated progress in clinical biomarker discovery, disease subtyping, and pathway-centric drug targeting.

Nevertheless, a persistent obstacle remains: how to bridge the gap between associative patterns in the data and the underlying functional organization of biological systems. Traditional approaches to pathway analysis can be broadly classified into statistical enrichment methods and machine learning frameworks[5]. Statistical enrichment tools typically identify pathways that are overrepresented among differentially expressed features, providing summary-level biological annotation lacking mechanistic detail [6, 7]. By contrast, recent advances in machine learning enable high-dimensional modeling of omics data with their pathway information, yielding powerful prediction models for clinical outcomes [8, 9]. However, such models treat molecular features as independent variables with limited capacity to capture the functional dependencies inherent to biological networks—and tend to operate as “black boxes” whose internal representations are difficult to interpret in biological terms [10].

An emerging paradigm in computational biology seeks to reconcile predictive performance with mechanistic explainability by integrating prior biological knowledge—such as pathway topology—directly into the architecture of machine learning models [8]. However, most strategies have so far relied on static graph embeddings or post hoc enrichment analysis, and rarely address the need to model the functional strength of molecular interactions. From a biological perspective, the effect a gene exerts on another, or the regulatory impact of a metabolite along a pathway, is fundamentally a functional mapping—one which varies in strength, directionality, and context [11, 12].

In this work, we present a knowledge-driven, explainable machine learning framework that elevates pathway analysis from conventional association testing to rigorous functional approximation. Inspired by the universal approximation theorem [13], we construct a neural network whose connectivity mirrors curated biological pathway graphs. Each node corresponds to a molecular feature, and each edge reflects a documented biological interaction, ensuring that the model structure is biologically plausible and interpretable. Crucially, our framework learns explicit, parameterized functional approximators for each interaction, represented as trainable mappings that quantify how one molecule influences another along the pathway. This formulation naturally accommodates the propagation of signals—analogous to biological processes such as gene regulation, signal transduction, and metabolic control—by leveraging residual connections that encode both direct and context-dependent effects. As a result, each model weight serves as a candidate functional coefficient, offering a bridge between statistical learning and mechanistic hypothesis generation.

Overall, this work positions functional approximation as a principled and generalizable methodology for mechanistic pathway analysis, unifying biological knowledge graphs, multi-omics integration, and explainable machine learning. Our framework catalyzes deeper insight into complex disease mechanisms, setting the stage for biologically grounded precision medicine.

## 2 Results

### 2.1 Method Overview

We propose a knowledge-driven machine learning framework for analyzing and interpreting omics data. The first core component is to integrate selected biomarkers with their corresponding biological pathways (Figure 1). The workflow begins with data acquisition and feature selection (Figure 1a). Numerous studies have explored approaches for selecting features or biomarkers [14–16], which we regard as crucial drivers of biological processes. Our framework leverages these selected features and is agnostic to the specific feature-selection method, provided that the features can be mapped to reference pathways. Subsequently, we construct a regulatory network over the selected features using curated pathway databases.

**Fig. 1.**
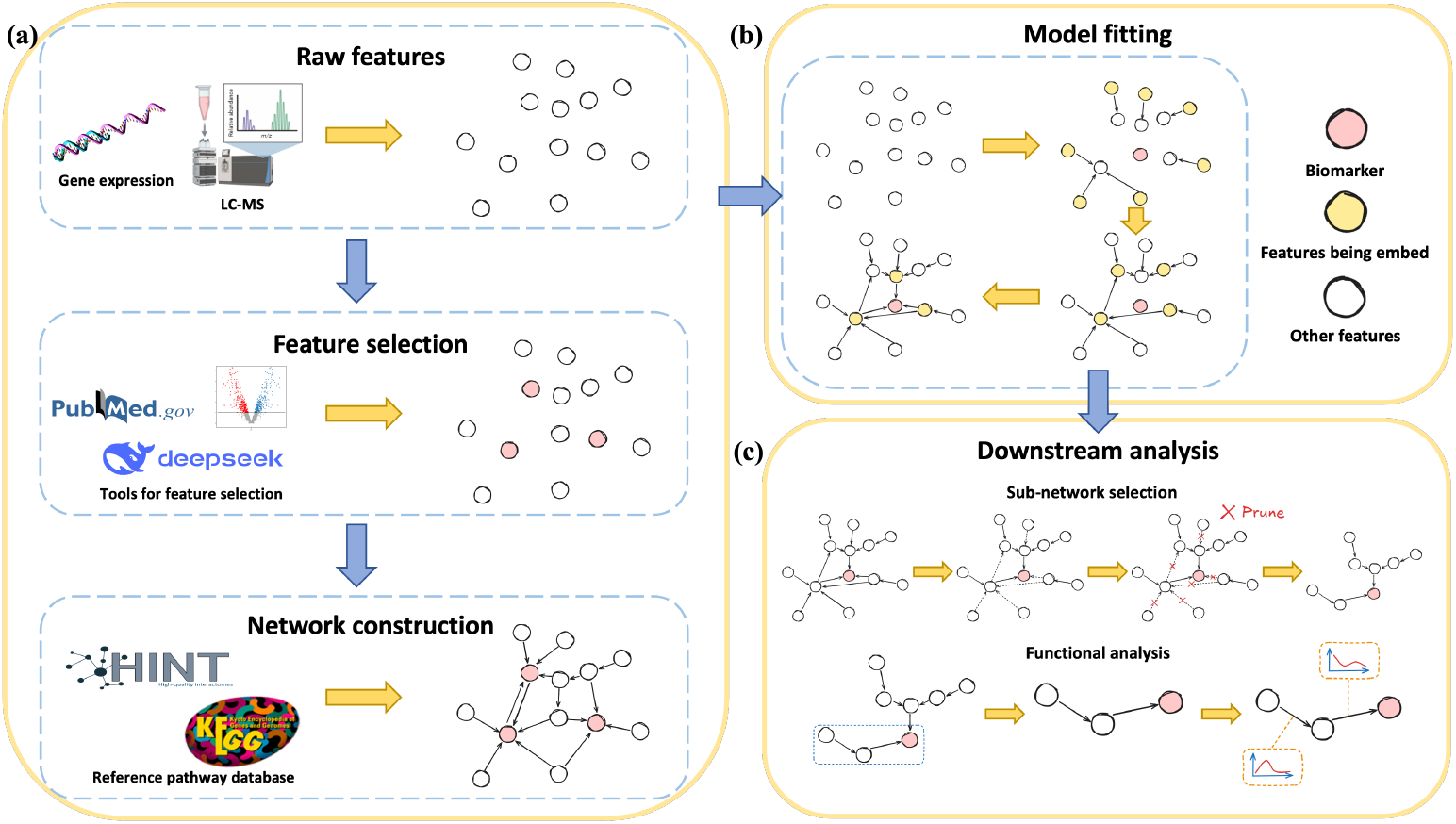
Overview of the whole method. **(a)** Procedures of pre-processing, including data acquisition like gene sequencing, LC-MS data, etc., feature selection using various tools and network construction with available databases. **(b)** A picture mimicking the model fitting procedure. Signals are modeled from marginal features to selected biomarkers. **(c)** Possible applications of our method, including select informative sub-network and analysis of the mutual relationship between biological molecular.

The second core component of our approach is a machine learning model that employs a network-like architecture (Figure 1b). The design is biologically inspired, mimicking interpathway signaling mechanisms in living organisms. In this knowledge-driven neural network, each neuron represents a specific biological feature. Connections between neurons are established only when the corresponding features participate in documented biological reactions. At each layer, we model how predetermined marginal features transmit signals toward selected key features. The cumulative effect of these marginal vertices propagates through the network to the key features, thereby characterizing pathway influences on the biomarkers. Finally, the adjusted values of the key features pass through a multilayer perceptron with softmax activation to assess their contributions to clinical outcomes.

The model is implemented in PyTorch [17] and trained with the Adam optimizer [18]. All experiments are conducted on a workstation with dual Intel Xeon Platinum 8276L CPUs (112 threads) and dual NVIDIA RTX 3090 GPUs (24 GB each).

### 2.2 Application on Breast Cancer data

MicroRNAs (miRNAs) are small non-coding RNA molecules that regulate gene expression post-transcriptionally through the miRNA-induced silencing complex (miRISC), where the mature miRNA guides the complex to complementary target mRNAs, resulting in translational repression or mRNA degradation [19, 20] (Figure 2b and c). For example, miR-373 has been shown to activate gene expression by targeting promoter sequences in DNA[21], underscoring the diverse regulatory roles of miRNAs beyond conventional mRNA targeting.

**Fig. 2.**
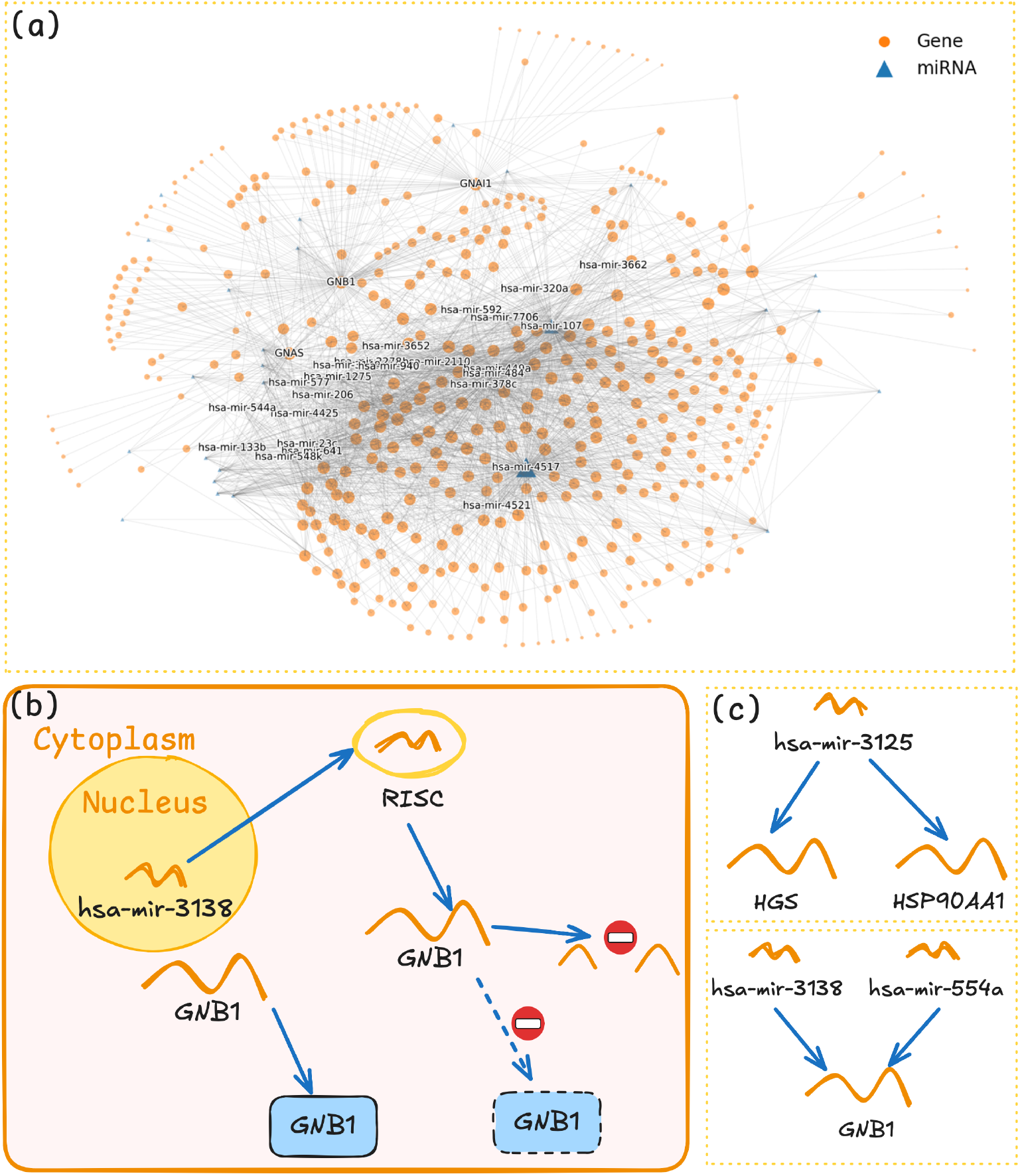
Overview of application on breast cancer dataset. **(a)** A draft of the selected gene regulatory network that is used in the experiment. Larger nodes indicates higher degrees of features in the graph. Nodes with degree ≥ 30 are labeled. **(b)** A draft of how miRNA function through RISC on gene expression. By binding with RISC, miRNA *hsa-mir-3138* resolves mRNAs and prevents the translation of the *GNB1* protein. **(c)** One type of miRNA can target multiple genes and a certain gene can be impacted by multiple miRNAs.

To investigate the influence of miRNAs on specific clinical outcomes mediated by gene regulation, we applied our method to the Cancer Genome Atlas (TCGA) breast cancer (BRCA) dataset[2]. For this analysis, we restricted our attention to matched miRNA expression (miRNA-Seq) and gene expression (RNA-Seq) profiles, thereby enabling a focused assessment of post-transcriptional regulatory interactions relevant to disease phenotype. We examined the association between miRNA/gene expression and a clinically relevant molecular subtype of breast cancer—estrogen receptor (ER) status, which plays a pivotal role in tumor progression [22].Understanding the biological mechanisms by which miRNAs and genes function within regulatory networks in association with ER status is of significant clinical importance. A key advantage of our method is its ability to integrate diverse molecular interactions, provided the relationship between any two molecular features is available. For gene-gene interactions, we referenced the HINT database [23], and for miRNA-target gene interactions, we used multiMiR [24] to match miRNAs with their gene targets.

#### 2.2.1 Identification of Genes Strongly Regulated by miRNAs

After filtering out genes and miRNAs with low expression values or lacking valid matching relationships, we retained 8,523 genes and 334 miRNAs for the analysis. The model was trained with a batch size of 16 for 100 epochs. In this experiment, we set the maximum network depth—excluding the effects of miRNAs—to three, and selected top-ranked features using a naive differential analysis, defined here as ranking features by their univariate statistical significance between case and control groups without applying multivariate modeling or extensive corrections. Such approaches are commonly used as a baseline in gene expression studies, where simple *t*-tests or fold-change criteria serve as initial filters for candidate features [25–27]. The final model achieved an average test accuracy of 0.952. A partial draft of the selected gene regulatory network is shown in Figure 2a.

Applying a stringent pruning threshold to the model enabled the identification of genes most strongly regulated by miRNAs, with potential relevance to estrogen receptor (ER) positivity. Although our framework may not always capture precise functional relationships, it robustly prioritizes genes that are under substantial miRNA regulatory influence and may play a role in key biological processes. Notably, many of these genes have previously been implicated in ER signaling or tumor progression, lending credibility to the selected candidates.

For example, studies have demonstrated that phytoestrogens such as genistein upregulate GNB1 expression through ER*α*/ER*β*-dependent transcriptional activation, directly linking ER signaling with downstream G-protein modulation [28, 29]. Likewise, BRD4 has been shown to potentiate ER activity via binding to acetylated histones, whereas disruption of SWI/SNF components leads to increased BRD4 recruitment and sustained ER-driven proliferation, even under anti-estrogen therapy [30].

In addition to pairs with previously established roles in ER signaling, our analysis also prioritized several miRNA–gene associations for which no direct evidence is currently available in the literature. Among the top miRNA–gene pairs identified, both HGS (targeted by hsa-mir-3125) and HSP90AA1 (also targeted by hsa-mir-3125) emerged as notable candidates, despite the absence of direct literature linking them to ER status in breast cancer. HGS (hepatocyte growth factor-regulated tyrosine kinase substrate) encodes a protein involved in endosomal trafficking and the regulation of receptor tyrosine kinase (RTK) signaling. While its role in modulating EGFR degradation and signal attenuation has been described in other malignancies, its potential involvement in hormone receptor pathways remains uncharacterized [31]. The identification of HGS in our analysis suggests that miRNA-mediated regulation of vesicular transport and RTK turnover could have as-yet unexplored implications for ER signaling or response to therapy.

Similarly, HSP90AA1 encodes a cytosolic heat shock protein 90 isoform, which functions as a molecular chaperone stabilizing a variety of client proteins, including many oncoproteins and kinases. HSP90 inhibitors have been investigated as potential therapeutics in several cancers, including breast cancer, largely due to their broad impact on oncogenic signaling networks [32, 33]. However, there is limited direct evidence associating HSP90AA1 with ER status specifically. The emergence of HSP90AA1 as a prioritized target in our miRNA–gene network points to the possible importance of post-transcriptional regulation of protein homeostasis in ER-positive tumors, warranting further functional investigation.

#### 2.2.2 Functional Interpretation

Beyond the immediate effects of miRNA regulation, several of the identified genes possess known roles in tumor biology and gene–gene regulatory networks. For instance, FETUB has been characterized as a tumor suppressor in prostate cancer, acting through inhibition of the PI3K/AKT pathway and modulation of apoptotic processes[34]. FETUB may also exert inhibitory effects in estrogen-driven contexts, and its interaction with meprin *α* (MEP1A) further implicates it in extracellular matrix remodeling and inflammation [35–38].

FoxG1 is another notable candidate, functioning as a tumor suppressor by disrupting the AIB1-E2F1-Sp1-p300 transcriptional activation complex at the AIB1 promoter[39, 40].Although FoxG1 does not directly interact with the estrogen receptor, its repression of AIB1—a critical ER coactivator—compromises ER-mediated transcription and impacts breast cancer progression and treatment resistance [41, 42]. The clinical association between low FoxG1 expression and poor prognosis underscores the relevance of this axis.

While the precise mechanistic cascades connecting these genes to ER status cannot be definitively established in this analysis, the literature support for their involvement in ER-associated pathways underscores the utility of knowledge-driven machine learning for hypothesis generation in molecular oncology. Further experimental validation and multi-omics integration are warranted to elucidate the underlying biological mechanisms.

### 2.3 Application on COVID-19 Metabolite Data

The robustness and flexibility of our knowledge-driven framework were further evaluated through its application to liquid chromatography - mass spectrometry (LC-MS) data in the context of COVID-19. LC-MS is a powerful analytical platform widely utilized in untargeted metabolomics to comprehensively profile metabolites within biological samples [43]. However, the inherent complexity of LC-MS data presents several analytical challenges, including signal drift, matrix effects, and the difficulties of metabolite identification given the vast chemical diversity and dynamic concentration ranges [44]. Moreover, preprocessing steps such as peak alignment, noise filtering, and normalization must be rigorously optimized to ensure data accuracy and reproducibility [45]. Addressing these challenges is critical for advancing untargeted metabolomics and enhancing biomarker discovery in disease research.

In this study, we leveraged our knowledge-driven modeling approach to analyze the ST001849 COVID-19 metabolomics dataset [46], which comprises LC-MS profiles from 609 test units collected over multiple time points and annotated with subsequent patient admission to the intensive care unit (ICU). The raw LC-MS data were obtained from the Metabolomics Workbench and processed using apLCMS [47], followed by batch effect correction with Combat [48]. A total of 913 metabolites were matched and integrated into the constructed pathway network, using the KEGG metabolic pathways as a reference [49]. This case study highlights the potential of our framework to interpret high-dimensional metabolomics data and elucidate clinically relevant molecular features in COVID-19.

#### 2.3.1 LLM-enhanced feature selection

Recent advances in large language models (LLMs) have enabled the integration of domain knowledge into computational workflows across biomedical research [50, 51]. In this study, we employed DeepSeek, a state-of-the-art LLM, to augment the feature selection process by prioritizing metabolites with potential clinical relevance to severe COVID-19 outcomes. The LLM-driven approach distills instructive information from existing biochemical databases and literature, facilitating the identification of functionally important metabolites from high-dimensional LC-MS data and streamlining the analytical pipeline.

After filtering, the selected metabolites associated with severe COVID-19 symptoms are listed in Table 1. The adoption of LLM-assisted feature selection not only improves interpretability, but also offers a pragmatic solution for extracting actionable biological knowledge from complex omics datasets. For instance, our analysis highlighted *β*-hydroxybutyric acid (BHB) as a top candidate. BHB, a ketone body produced under metabolic stress, has recently been shown to modulate immune responses by inhibiting the NLRP3 inflammasome—a central driver of cytokine storms in severe COVID-19 [52]. Furthermore, BHB may exert antiviral effects by altering cellular metabolism and restricting viral replication [53].

**Table 1.** LLM Selected Metabolites with Their KEGG ID

| KEGG ID | Name |
| --- | --- |
| C00328 | L-Kynurenine |
| C00186 | (S)-Lactate |
| C00584 | Prostaglandin E2 |
| C02165 | Leukotriene B4 |
| C00696 | Prostaglandin D2 |
| C01089 | $\beta$ hydroxybutyric acid |
| C00042 | Succinate |
| C06124 | Sphingosine 1-phosphate |
| C00249 | Hexadecanoic acid |
| C00780 | Serotonin |

#### 2.3.2 Metabolic subnetwork associated with COVID progression

The model was trained with a batch size of 16, a maximum depth of 3, and for 100 epochs. The average test accuracy achieved was 0.725. Among the selected metabolites and pathways, the tryptophan metabolic pathway (see Figure 3) was particularly prominent. Dysregulation of tryptophan metabolism has been implicated in the pathogenesis of severe COVID-19 and SARS, especially through its immunomodulatory effects. Tryptophan is metabolized via the kynurenine pathway, which is activated by pro-inflammatory cytokines such as interferon-gamma (IFN-*γ*), leading to the production of immunoregulatory metabolites such as kynurenine. Elevated kynurenine levels have been associated with immune suppression and increased disease severity in COVID-19, potentially exacerbating viral persistence and hyperinflammation [54]. In addition, prostaglandins—particularly prostaglandin E2 (PGE2)—play a central role in modulating inflammatory responses during viral infections. Overproduction of PGE2 has been linked to cytokine storms and acute respiratory distress syndrome (ARDS) in severe COVID-19, as it enhances vascular permeability and leukocyte recruitment [55]. Some reactions with in this metabolism are also shown in Figure 3. For example, KEGG compound C00028 (ATP) is the central energy currency of the cell. Its synthesis depends on both nitrogen and one-carbon metabolism: C00014 (ammonia) supplies nitrogen via glutamine and glutamate, while C00058 (formate) donates one-carbon units for purine biosynthesis [56]. Excess ammonia impairs oxidative phosphorylation and lowers ATP [57], and disrupted one-carbon flux can alter nucleotide pools. Although direct links are lacking, mitochondrial dysfunction and energy imbalance are consistently reported in severe COVID-19 [58]. Taken together, these metabolic perturbations highlight the therapeutic potential of targeting tryptophan and prostaglandin pathways to mitigate severe disease outcomes.

**Fig. 3.**
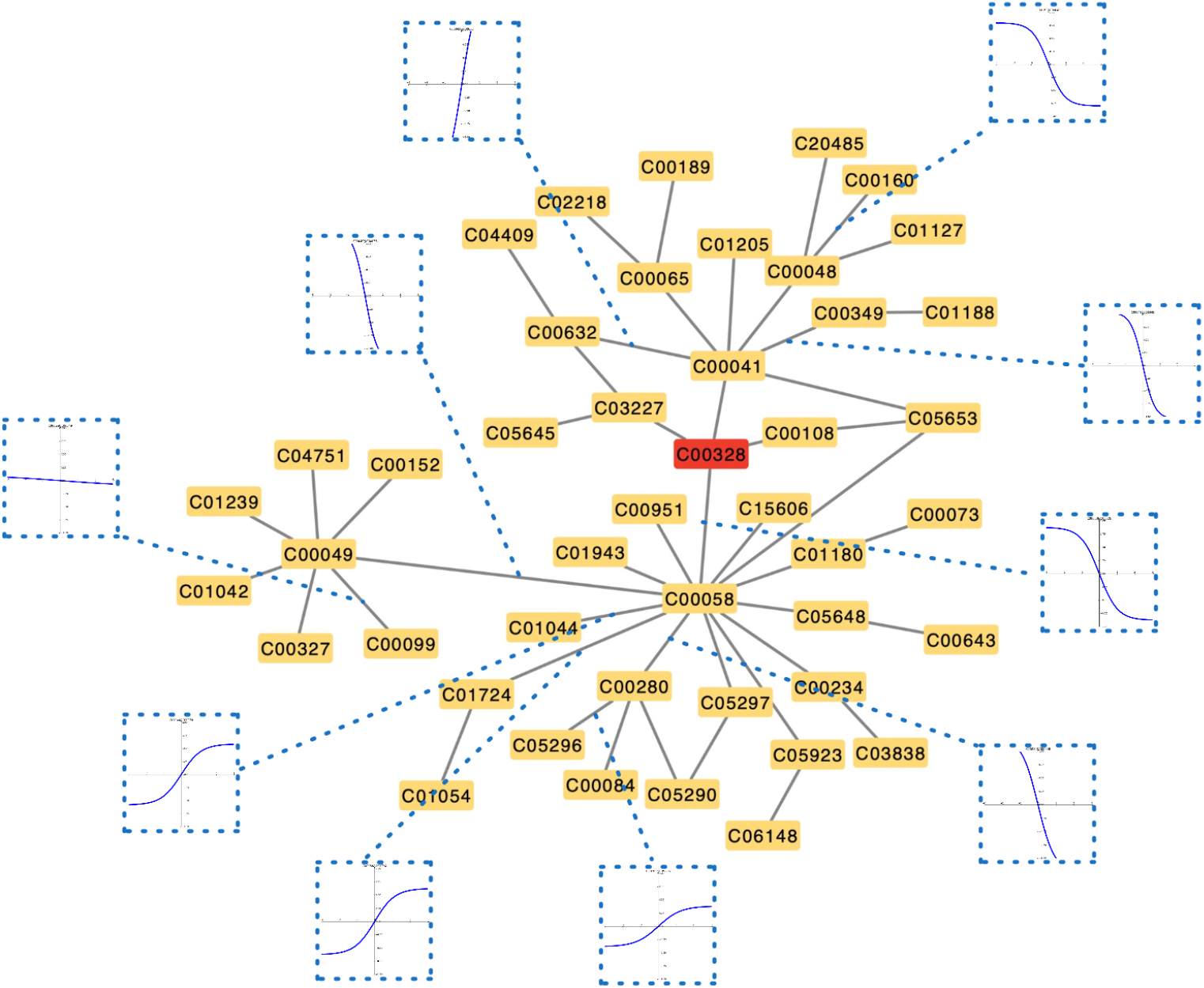
A topology graph of the selected tryptophan metabolic pathway. Each nodes represents a specific metabolite and an edge exists if two metabolite have certain reactions. Several edges’ function approximation are centralized and displayed on the same scale.

The arachidonic acid (AA) metabolism pathway, encompassing both prostaglandins and leukotrienes, was also selected and is known to play a pivotal role in the inflammatory response associated with severe COVID-19 and SARS infections. Following viral invasion, phospholipase A2 (PLA2) releases AA from membrane phospholipids, which is subsequently metabolized by cyclooxygenase (COX) and lipoxygenase (LOX) enzymes into prostaglandins (PGs) and leukotrienes (LTs), respectively. Prostaglandins, particularly PGE2, mediate fever, vasodilation, and pain, whereas leukotrienes (e.g., LTB4 and cysteinyl-LTs) promote neutrophil recruitment and bronchoconstriction, thereby exacerbating respiratory distress in severe cases [55, 59]. Evidence indicates that SARS-CoV-2 infection upregulates COX-2 expression, resulting in excessive prostaglandin production that may contribute to cytokine storms and acute lung injury [60]. Furthermore, leukotriene-driven inflammation has been implicated in pulmonary fibrosis, a long-term sequela of severe COVID-19 [61]. Accordingly, pharmacological inhibition of AA metabolites—such as through COX-2 inhibitors or leukotriene receptor antagonists—has been proposed as a potential therapeutic strategy to attenuate hyperinflammation in severe viral infections [61, 62]. Lactic acid is also selected as a biomarker, whose accumulation further exacerbates immune dysregulation by suppressing antiviral T-cell responses and promoting cytokine storms, a hallmark of severe COVID-19 [63]. Additionally, lactate has been shown to stabilize hypoxia-inducible factor 1-alpha (HIF-1*α*), which enhances viral replication by upregulating angiotensin-converting enzyme 2 (ACE2) expression, the primary entry receptor for SARS-CoV-2 [53]. These findings suggest that lactic acid not only serves as a biomarker of disease severity but also actively contributes to the progression of severe viral infections by modulating host metabolism and immune responses. However, no relevant metabolites that may interact with lactic acid is detected and matched in reference to our pathway, suggesting that further analysis is needed for structural function of metabolism.

Collectively, these results illustrate the value of combining LLM-guided feature selection and knowledge-driven machine learning for prioritizing biologically and clinically relevant metabolic sub-networks.

### 2.4 Simulation and ablation study

#### 2.4.1 Simulation Design

To evaluate the performance and interpretability of our model, we conducted simulation and ablation studies. For simplicity, we restricted the setting to single-omics data. The nodes generated in this simulation can be regarded as gene expression data.

There exists extensive work on graph generation, including scale-free networks such as the Barabási–Albert (BA) model [64], as well as studies dedicated to simulating gene expression data [65, 66]. However, our goal here was not to generate graphs with hubs of very high degree, nor to simulate fully realistic biological datasets. Instead, we reused the same reference network as in Section 2.2 and applied a relatively simple yet reasonable algorithm to generate gene expression count data suitable for model validation.

We first sampled nodes from the reference graph while retaining their original connections. To construct the feature abundance matrix, we assumed — without loss of generality — that the marginal distribution of each feature followed a Gaussian distribution with mean zero and variance one. Because gene expression data typically contain uninformative dimensions and dropout events, we randomly masked a fraction of entries to mimic sparsity.

We then simulated the effect of one feature on others along the reference pathway graph. Specifically, for a given feature *X*_*i*_ with value *x*_*i*_, any node *X*_*j*_ within distance *d* was perturbed by

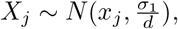

where *σ*_1_ is the impact coefficient. We assumed no accumulative effect; i.e., the perturbation of one node did not propagate recursively to its neighbors. Based on this procedure, we constructed *X*_before_ and *X*_after_ ∈ R^*n×p*^, and assigned labels according to values of deterministic features in *X*_after_.

Among features with degree above a chosen threshold, we subsampled nodes as informative features, since high-degree nodes are more strongly affected by their neighbors even with relatively large maximum step lengths. This strategy adds noise and difficulty to the prediction task, thereby providing a rigorous test of our method. After selecting informative nodes, we first simulated the impact of all other nodes on them, and then used a subset of these to determine labels. For deterministic parameters *w*, we sampled *w* ∼ *N* (0, *σ*_2_) and defined labels **y** ∈ {0, 1}^*n*^ by

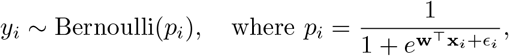

for each **x**_*i*_ ∈ *X*_after_, with *ϵ*_*i*_ denoting independent random noise.

#### 2.4.2 Simulation Details

To further assess scalability, we varied the dimensionality of the simulated datasets. The feature size was set to 1000, 2000, 4000, and 8000. In cases where the number of samples *n* was smaller than the number of features *p*, we selected sample sizes equal to 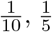, and 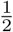 of the feature size. Among all features, 1% were designated as informative and 5% were randomly up- or down-regulated.

We assumed perfect modeling of relationships within the reference graph. That is, the impact radius *d* was set equal to *γ*_1_, the maximum depth of signal propagation. Because simulated data can be unstable across different realizations, for each fixed combination of sample size and feature size we repeated the simulation ten times. To maintain balanced classes, we constrained the proportion of samples in either class between 45% and 55%.

Models were then trained on these datasets with hyperparameters varied systematically. We compared our knowledge-driven framework against standard baselines including logistic regression, random forest, and feed-forward neural networks. Performance was evaluated using accuracy on held-out test sets, and interpretability was measured by the ability to recover ground-truth informative subnetworks.

Results showed that our method consistently achieved competitive accuracy while outperforming baselines in interpretability (Figure 4). Specifically, the model effectively identified subnetworks corresponding to the designated informative features, whereas conventional methods either failed to recover these structures or relied heavily on feature selection heuristics. Ablation experiments confirmed that removing pathway knowledge substantially reduced interpretability, thereby demonstrating the value of embedding prior biological information into model architecture.

**Fig. 4.**
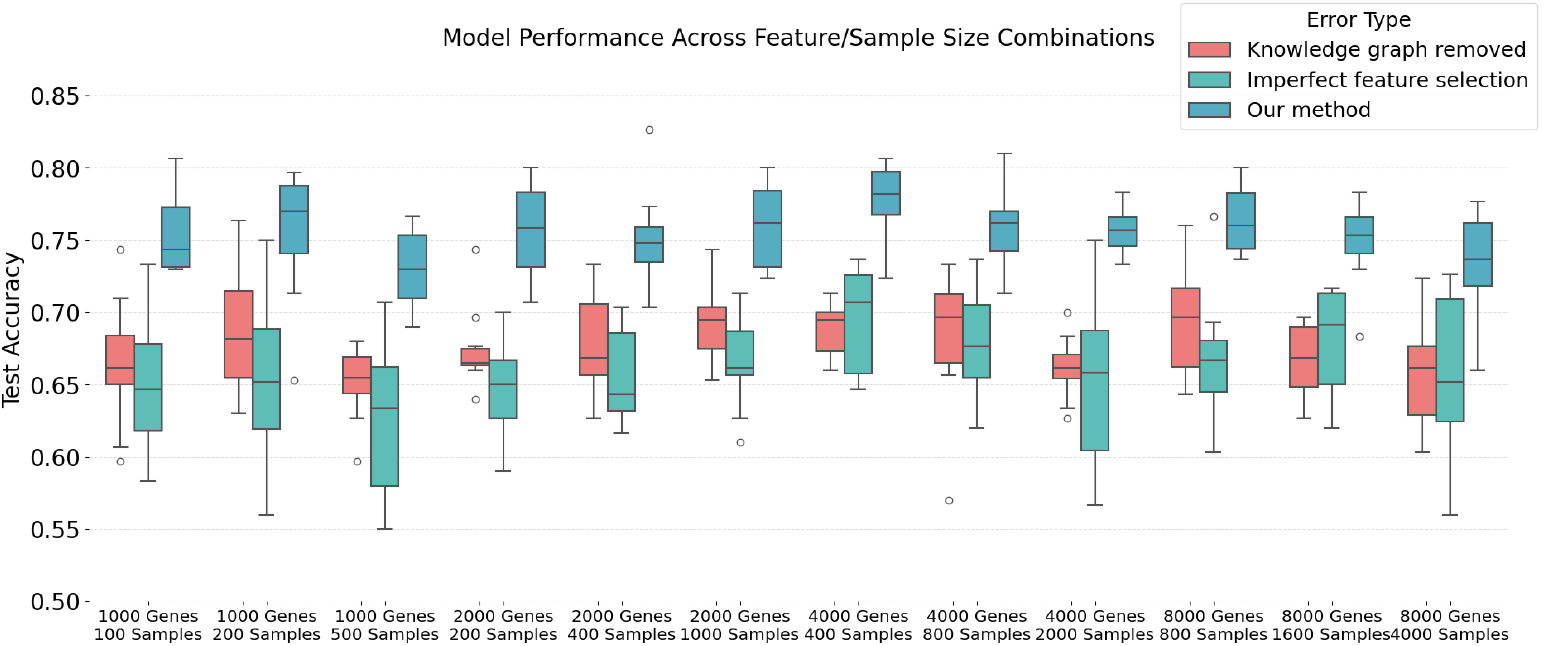
Prediction accuracy of our method. X-axis shows the combination of different feature size and sample size.

Overall, these simulation studies demonstrate the robustness of our framework under high-dimensional settings (*n < p*) and validate its capacity to balance predictive performance with interpretable pathway discovery.

## 3 Discussion

A key advantage of our approach is the natural integration of omics data with their underlying pathway structures. This design ensures that the architecture and downstream analyses follow biologically meaningful logic. Consequently, all edges in the neural network are interpretable, offering insights into mechanistic behavior rather than serving solely as predictive associations. To facilitate practical use, we have developed the package `PathNet`, which streamlines the application workflow and supports further exploration across diverse datasets.

Despite these strengths, several limitations remain. First, although our framework provides interpretable pathway-level insights, it cannot capture the full complexity of biological systems. Incomplete pathway annotation and context-dependent interactions restrict mechanistic resolution. Moreover, when modeling complex biochemical reactions among molecular features, embedding information solely from a pathway perspective yields only a partial understanding of reality and constrains interpretation of known reactions. Future work should consider probabilistic pathway modeling, as has been applied in graph-based analyses of spatial transcriptomics [67]. Such approaches would enable not only adjustment of pathway annotations but also the discovery of undocumented interactions.

Second, the reactions between features are simplified in our current implementation. In practice, the causal impact of one molecular feature on another is often directional. Ignoring such directionality may produce calculations and inferences that are difficult to justify or interpret. Incorporating directional connections into the model, however, introduces challenges: long reaction chains require deeper neural architectures, which increase model complexity and uncertainty of interpretation. Furthermore, the order in which each feature is embedded currently follows the intuitive distance between nodes on the topological graph. Future research should investigate more flexible embedding strategies that propagate marginal features toward central nodes while accounting for directionality and reaction hierarchy.

## 4 Conclusion

In this study, we proposed an explainable machine learning framework for analyzing omics data within a knowledge-driven pathway context. The model accommodates both homogeneous and heterogeneous data types, demonstrating substantial flexibility. To our knowledge, this is the first approach to simultaneously leverage graph-structured pathway information and model reactions among features within a unified framework. By embedding marginal features and propagating informative signals through biological processes, the framework provides a biologically intuitive representation. Applications to both the miRNA–gene breast cancer dataset and the COVID-19 LC-MS dataset highlight the broad applicability of our method.

## 5 Methods

### 5.1 Knowledge-driven Interpretable Neural Network

We introduce a novel neural network architecture based on a sparsely connected multilayer perceptron (MLP). By constraining the network connectivity according to curated biological pathways, the design mitigates overfitting while retaining strong predictive ability and interpretability [68]. The central idea is to emulate biological signal transmission: information from marginal molecular features flows along pathway connections toward a subset of biologically important centric features, which are ultimately used for outcome prediction.

Formally, let **X** ∈ ℝ^*n×p*^ denote the feature matrix, where *n* is the number of samples and *p* is the number of molecular features, and let *y* ∈ ℝ^*n*^ be the outcome labels. Prior biological knowledge is given in the form of a graph *G* = (*V, E*), where each vertex *v*_*i*_ ∈ *V* represents a distinct molecular feature (e.g., a gene or metabolite) and each undirected edge *e*_*ij*_ ∈ *E* corresponds to a known biological interaction derived from pathway databases.

Within *V*, we first select *γ*_2_ *centric features*, denoted 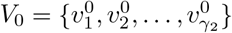, chosen for their potential biological relevance to the prediction task. The remaining features *V\V*_0_ are considered *marginal features* that may influence the centric features through pathway connections. To organize the network layers, we introduce the hyperparameter *γ*_1_, representing the maximum number of hops along *G* over which a marginal feature can transmit information to a centric feature.

For each non-centric feature *v*_*i*_ ∈ *V \ V*_0_, we compute its shortest-path distance to the nearest centric feature in *V*_0_, denoted

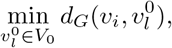

where *d*_*G*_(*·, ·*) is the standard graph distance in *G*. If this minimum distance equals *j* ≤ *γ*_1_, the feature *v*_*i*_ is assigned to the set *V*_*j*_. Thus, *V*_1_ contains features one hop away from *V*_0_, *V*_2_ contains those two hops away, and so on up to 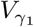. This assignment procedure is summarized in Algorithm 1 and illustrated in Figure 1(b).

The network then consists of *γ*_1_ layers. In layer *j*, features in *V*_*j*_ transmit updated signals to their immediate upstream neighbors in *V*_*j*−1_, progressively aggregating peripheral information toward the centric features in *V*_0_. After this stepwise aggregation, the updated centric feature representations are passed to a fully connected multilayer perceptron with softmax activation to produce the final prediction.

#### 5.1.1 Biology-Inspired Residual Connections with Learnable Functional Approximation

Residual, or shortcut, connections have been shown to preserve information flow and improve optimization stability by mitigating gradient vanishing in deep neural networks [69]. To emulate analogous information gain and signal propagation mechanisms in biological systems, we model the influence of one molecular feature *v*_*i*_ on another *v*_*j*_ as

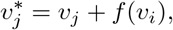

where *f* represents the functional impact of *v*_*i*_ on *v*_*j*_. In this formulation, the updated feature 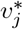 is a combination of its original value and the contribution from interacting features.

Inspired by the universal approximation theorem [13], we parameterize *f* as

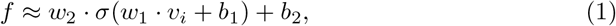

where *w*_1_, *w*_2_, *b*_1_, *b*_2_ are trainable parameters and *σ* is an element-wise activation function. This two-layer functional form is sufficiently expressive to capture nonlinear biological influences while remaining simple enough to reduce overfitting. Features without documented interactions maintain their values without modification.

To express this transmission process compactly, let 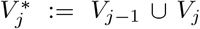 be the set of features in consecutive layers *l*_*j*_ and *l*_*j*+1_. Define the adjacency matrix 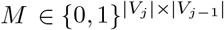 by

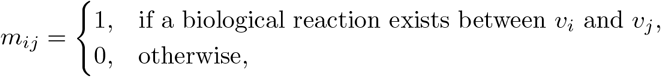

and the residual connection matrix 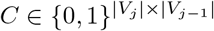 whose *i*th row **c**_*i*_ is

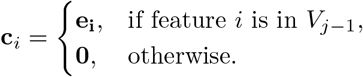

Introducing *C* enables the residual connection. For input 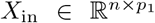, the transformation from *V*_*i*_ to *V*_*j*_ is:

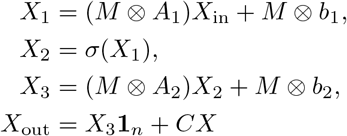

where 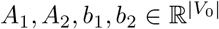 are trainable parameters, *σ* is the activation function, 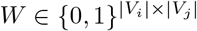 is the adjacency matrix, and ⊗ denotes element-wise multiplication. The resulting adjusted centric feature values 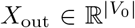 are then passed to a fully connected layer with softmax activation for outcome prediction.

### 5.2 Sub-network Selection and Functional Analysis

Extensive studies have explored knowledge distillation and sub-network selection in deep neural networks [70–72]. A simple yet effective strategy is to prune all weights whose absolute values fall below a threshold [73], which can preserve predictive performance while enhancing interpretability. In our framework, we apply a similar pruning principle to identify informative subnetworks within the overall pathway.

Specifically, in the functional approximation defined in Equation 1, the parameters *w*_1_ and *w*_2_ can be interpreted as the interaction strength between features *v*_*i*_ and *v*_*j*_. Thus, an intuitive method for sub-network selection is to remove edges whose *w*_1_ values are small in magnitude. In such cases, the signal transmission effectively reduces to an identity mapping, and the source feature *v*_*i*_ can be regarded as uninformative (Figure 1c).

Previous work on graph-structured machine learning has used connection weights to quantify the importance of features [74–76]. In many domains, such as computer vision, interpreting the meaning of a connection between two neurons can be difficult [77, 78]. In contrast, a key advantage of our approach is that each edge corresponds to a documented molecular reaction from a knowledge graph or curated database. The functional estimator in Equation 1 therefore provides both an approximation of the underlying biological interaction and a natural mechanism for regularization to avoid overfitting.

After pruning edges with low absolute weights, the remaining connections can be visualized to reflect strong functional relationships between molecular features, with large slopes indicating greater relevance to the outcome of interest. For completeness of pathway interpretation, features not directly analyzed can be reintroduced into the selected sub-network as context nodes, ensuring biological coherence.

## Declarations

### Data availability

The data analyzed in this study originated from public data. The longitudinal metabolomics of COVID-19 disease severity data is downloaded from metabolomics workbench(https://www.metabolomicsworkbench.org/data/DRCCMetadata.php?Mode=Study&StudyID=ST001849). The gene and miRNA expression can be found on NIH gdc data portal(https://portal.gdc.cancer.gov). We have also provided sample dataset that has been filtered in our code repository.

### Code availability

To assist further exploration and validation of our method, we have developed a package *PathNet*, which is available on https://github.com/YanLarryKE/PathNet.

## Acknowledgments

This work was partially supported by the National Key R&D Program of China (2022ZD0116004), Guangdong Talent Program (2021CX02Y145), Guangdong Provincial Key Laboratory of Big Data Computing, and Shenzhen Science and Technology Program ZDSYS20230626091302006 and JCYJ20240813113536047.

## Appendix A Algorithm for model construction

### Algorithm 1

Feature vertices layer partitioning

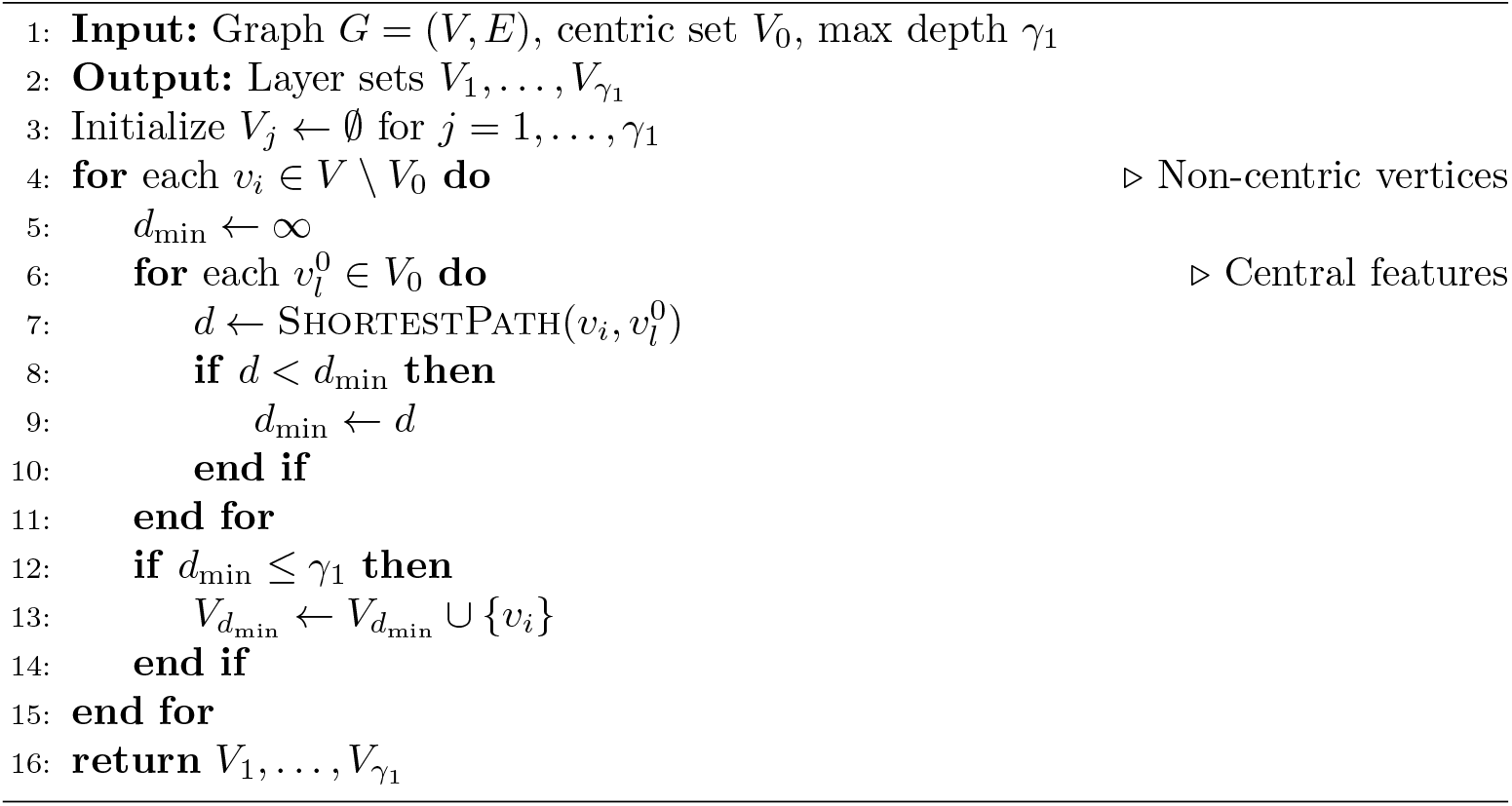

